# Metabolome shift associated with thermal stress in coral holobionts

**DOI:** 10.1101/2020.06.04.134619

**Authors:** Amanda Williams, Eric N. Chiles, Dennis Conetta, Jananan S. Pathmanathan, Phillip A. Cleves, Hollie M. Putnam, Xiaoyang Su, Debashish Bhattacharya

## Abstract

Coral reef systems are under global threat due to warming and acidifying oceans^1^. Understanding the response of the coral holobiont to environmental change is crucial to aid conservation efforts. The most pressing problem is “coral bleaching”, usually precipitated by prolonged thermal stress that disrupts the algal symbiosis sustaining the holobiont^2,3^. We used metabolomics to understand how the coral holobiont metabolome responds to heat stress with the goal of identifying diagnostic markers prior to bleaching onset. We studied the heat tolerant *Montipora capitata* and heat sensitive *Pocillopora acuta* coral species from the Hawaiian reef system in Kāne’ohe Bay, O’ahu. Untargeted LC-MS analysis uncovered both known and novel metabolites that accumulate during heat stress. Among those showing the highest differential accumulation were a variety of co-regulated dipeptides present in both species. The structures of four of these compounds were determined (Arginine-Glutamine, Lysine-Glutamine, Arginine-Valine, and Arginine-Alanine). These dipeptides also showed differential accumulation in symbiotic and aposymbiotic (alga free) individuals of the sea anemone model *Aiptasia*^4^, suggesting their animal provenance and algal symbiont related function. Our results identify a suite of metabolites associated with thermal stress that can be used to diagnose coral health in wild samples.

---

The exchange of metabolites, either between organism-environment or organism-organism, gave rise not only to early life, but complex life systems on Earth. Examples include bacterial deep vent communities^5^, the plant rhizosphere^6^, the human microbiome^7^, and the coral holobiont^8^. Exchange of metabolites between stony corals (Scleractinia), their dinoflagellate algal photosymbionts (Symbiodiniaceae), and associated microbes is the foundation for modern coral reef ecosystems^1,9^ that cover ca. 255,000 km^2^ of the planet surface^10^. Under ambient conditions, the algal cells provide 90-95% of host energy needs in the form of lipids, carbohydrates, amino acids, and O_2_^11^. In return, excess nitrogen and inorganic waste from the coral animal are recycled by the algae to fuel cell metabolism^12,13^. Environmental stress disrupts this symbiotic relationship, resulting in coral bleaching^2^. During this stressful process, coral susceptibility to disease increases^14^ as growth slows and reproduction rate decreases^15,16^. However, the holobiont is the fundamental unit of selection with the host cnidarian playing the dominant role in recovery and resilience^17^. Here we used metabolomics with *M. capitata* and *P. acuta* (Fig. 1a) coral holobionts to connect biochemistry to physiology to ecology^18,19^ by identifying metabolic features associated with the short-term thermal stress response. Previous work demonstrates that Hawaiian *M. capitata* can meet 100% of its daily metabolic energy requirements through heterotrophic feeding during periods of bleaching^17^, whereas *P. acuta* from this reef system experiences increased thermal sensitivity^20^. Nubbins from these two coral species were exposed for 5-weeks (T1-T5) to a 2.7**°-** 3.2**°**C temperature increase and sampled at four time points (see Methods). Three time points (T1, T3, T5) represent tank sampling, whereas T0 nubbins were collected from local wild populations (Fig. 1b). During the experiments, Kāne’ohe Bay was at the beginning of an unexpected warming event of ca. 1**°**C, leading to thermal stress in the ambient condition tanks that drew water from the Bay and subsequently a bleaching event that peaked in September-October 2019. The impact of this natural fluctuation was recorded in photographic bleaching scores that measure color as a proxy for algal abundance in the coral nubbins^21^ (Fig. S1). *M. capitata* showed little visible effect from the warming event, whereas *P. acuta* in the ambient tanks was impacted in a similar fashion as the high temperature treatment.

**Fig. 1.**
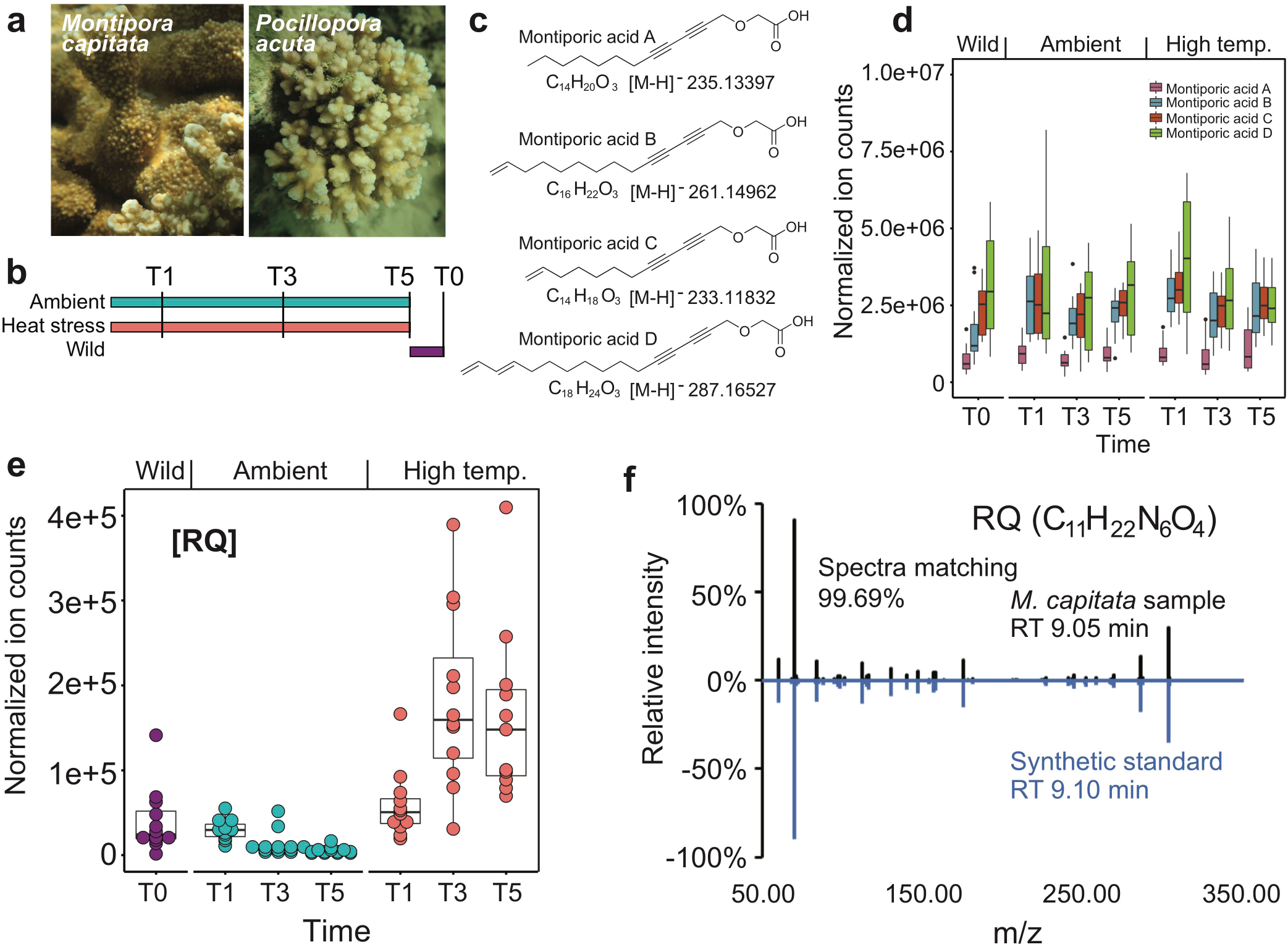
Analysis of Hawaiian stony corals. (**a**) Images of *M. capitata* and *P. acuta* from Kāne’ohe Bay, O’ahu. (**b**) Experimental plan. (**c**) Structures of Montiporic acids identified in the coral holobiont. (**d**) Accumulation of total MAs in *M. capitata* over the duration of the tank experiments as well as from wild populations. (**e)** Metabolite RQ in *M. capitata* accumulated over time during thermal stress (T1-T5). (**f**) The metabolite C_11_H_22_N_6_O_4_ that showed accumulation under heat stress matches synthetic standard of RQ dipeptide in retention time and MS^2^ spectra.

Untargeted LC-MS metabolomics revealed thousands of features from the coral samples, many of which showed a significant difference in accumulation under ambient and high temperature treatments (Fig. S2a). Characterized metabolic features included central metabolites such as amino acids, nucleotides, and sugar phosphates. In addition, high-resolution LC-MS^2^ allowed the detection, quantitation, and structural elucidation of many secondary metabolites. An example is Montiporic acids (MAs) (Fig. S2b), which were first discovered in Australian coral eggs^22^. These compounds are disubstituted acetylenes with a carboxyl group linked to two alkyne (carbon-carbon triple bond) groups, followed by an unbranched alkane / alkene tail. The four known MAs (MA-A – MA-D; Fig. 1c) have antimicrobial activity, cytotoxicity against leukemia cells, and reduce the photosynthetic competency of coral symbionts^22,23^. The MAs are highly abundant in the *M. capitata* samples from both ambient and heat stressed conditions as well as in the wild samples (Fig. 1d), exceeding by >1,000-fold the amounts in wild *P. acuta* samples (Fig. S2c). This suggests that MA metabolic activity in different coral species can vary significantly. It is unclear whether MAs contribute to stress resistance in *M. capitata* by controlling symbiont density prior to warming events (see below). Nonetheless, the high abundance of MAs in *M. capitata* suggests coral animals divert a large portion of metabolic resources to the production of secondary metabolites.

We were interested in other known and novel and known metabolomic features in the holobiont that accumulate under high temperature treatment and may diagnose thermal stress prior to visible bleaching. By comparing the metabolomic features of the ambient and heat stressed T1-T5 groups in *M. capitata*, we observed many features that have significantly increased abundance in the heat stressed group. One feature, m/z 303.177 at retention time 9.05 min, was detected under positive ionization mode in *M. capitata* and showed the highest accumulation over time. This feature also showed a time-dependent increase under high-temperature in *P. acuta* (Fig. S3a). Based on the accurate mass and isotopic fine structure, this molecule was assigned the chemical formula of C_11_H_22_N_6_O_4_. To gain insights into the structure of this metabolite, we collected the MS^2^ spectra using parallel reaction monitoring (PRM) and generated high quality MS^2^ pseudospectra by calculating the correlation between MS^1^ and MS^2^ spectra. Through systematic comparisons of the accurate mass, retention time and the MS^2^ spectra for chemically synthesized pure standards of various dipeptides and tripeptides (Table S1), we positively identified the C_11_H_22_N_6_O_4_ to be Arginine-Glutamine (RQ; Figs. 1e, 1f).

The accumulation of RQ dipeptide under heat stress could result from increased proteolysis and/ or insufficient peptide clearance. We speculate that both factors contribute to the elevation of not just RQ but also other dipeptides. This idea is supported by the untargeted metabolomics results. Among the metabolites that show significantly different levels between ambient and heat stressed conditions, the majority match the accurate mass of dipeptides (Figs. 2a-c). These putative dipeptides are significantly enriched under the heat stressed condition at both T3 (*p* = 4.8e-13, Fisher’s Exact Test) and T5 (*p* = 5.6e-14) time points. To confirm the chemical identities of these putative dipeptides, we synthesized the chemical standards and collected MS^2^ spectra from the coral samples and the standards. The structures of three additional metabolites were determined: C_11_H_22_N_4_O_4_ ([M+H]^+^ m/z 275.1714 [Lysine-Glutamine, KQ], Figs. 2d, 2g), C_11_H_23_N_5_O_3_ (m/z 274.1874 [Arginine-Valine, RV; Figs. 2e, 2h]), C_9_H_19_N_5_O_3_ (m/z 246.1561 [Arginine-Alanine, RA; Figs. 2f, 2i]). Based on these results, we conclude that dipeptides are important markers of stress in *M. capitata*. It should be noted that some features not identified as dipeptides also increased in intensity with thermal stress, however, the majority are not statistically significant (Fig. 2c). In comparison, the stress sensitive and bleached *P. acuta* showed many metabolites with decreased abundance after heat stress, reflecting a dampened metabolism. However, the RQ and KQ dipeptides showed significant accumulation in this coral at T3 and T5 (Fig. S3a). The similarities and clear differences in dipeptide production in these two coral species suggest that their production may not be explained solely by proteolysis due to metabolic shutdown prior to cell death and that they are bona fide early metabolic signals of bleaching in response to thermal stress. We did not determine which component of the holobiont produced each metabolite, however, an accumulation of proteogenic dipeptides in *Arabidopsis thaliana* during a time-course stress experiment is linked to autophagy^24^. Dipeptides also serve a diversity of other functions such as small molecule regulators (e.g., H^+^ buffers^25^, antioxidants^26^, and glucose regulators^27^). Specifically, RQ can ameliorate the impacts of oxygen imbalance in a murine model of retinopathy of prematurity (ROP^28^) and may have beneficial effects on the reversal of oxygen-induced lung damage^29^. Therefore, we suggest that RQ may be a response to oxygen stress resulting from redox imbalance in coral algal symbionts caused by thermal stress.

**Fig. 2.**
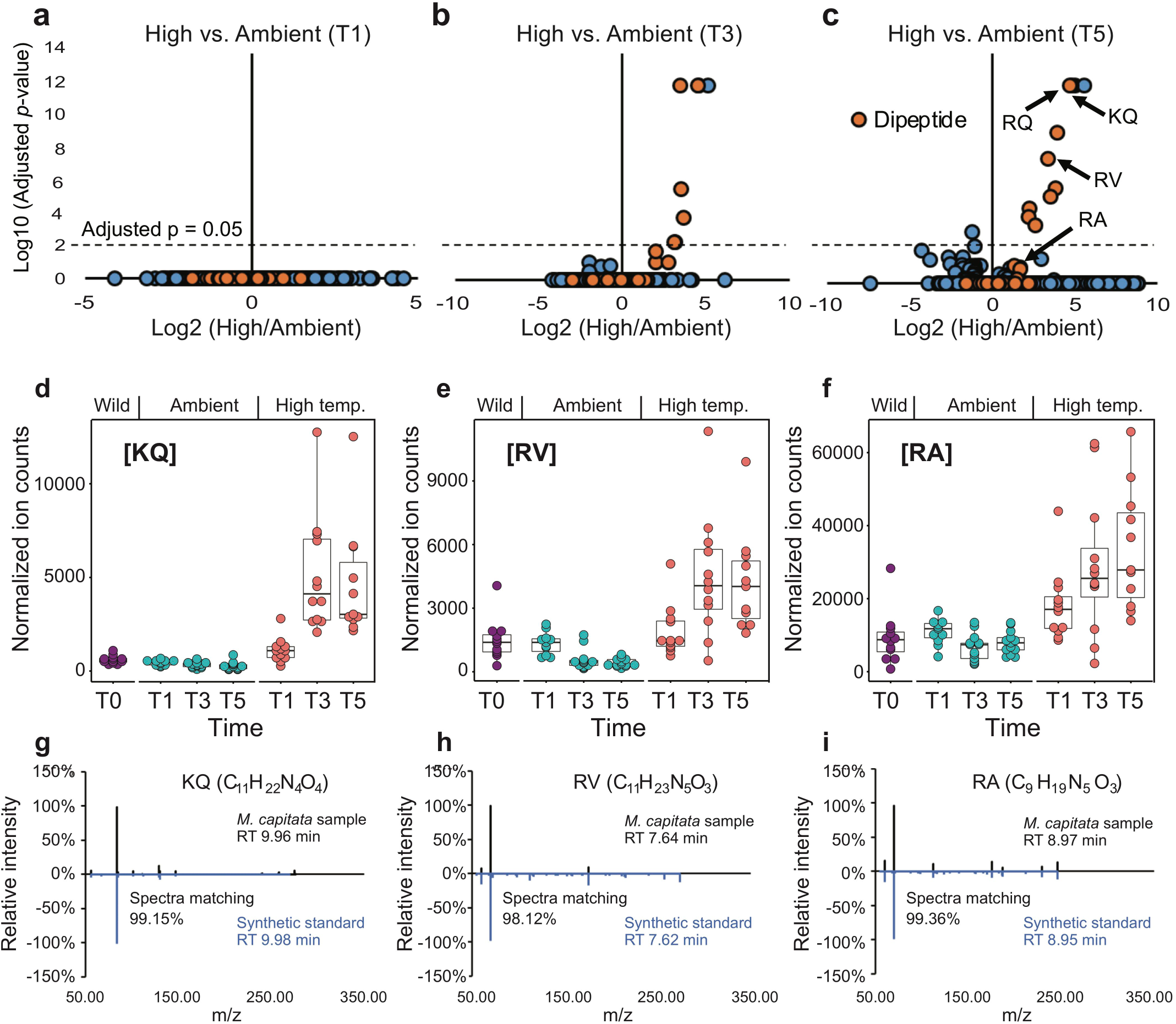
Dipeptide production by stony corals under thermal stress. (**a-c**) Volcano plots of *M. capitata* metabolites generated from positive mode data when comparing ambient and high temperature treatments. Putative dipeptides are shown with the filled orange circles. **(d-f)** Accumulation of dipeptides KQ, RV, and RA under heat stress. (**g-i**) The metabolites that showed accumulation under heat stress matche synthetic standards of KQ, RV and RA dipeptides in retention time and MS^2^ spectra.

Targeted analysis showed that some known primary metabolites also accumulate during prolonged thermal stress in *M. capitata*. Under high temperature, proline, methionine, methionine sulfoxide, and CDP-choline are all significantly increased at T5 (Figs. 3a-d). The accumulation of methionine is of particular interest because of its involvement in scavenging reactive oxygen species (ROS) and initiating and maintaining epigenetic modifications. Methionine and methionine sulfoxide are both upregulated in response to prolonged heat stress, whereas S-adenosyl-L-methionine (SAM) is downregulated. The increase of methionine and methionine sulfoxide is likely due to the role of methionine in scavenging ROS^30,31^, with ROS generated from the symbiotic breakdown^32^ driving methionine oxidation that increases methionine sulfoxide levels. Further, apoptotic and necrotic pathways triggered by thermal stress would likely release additional methionine from cellular pools. The reduction of SAM availability may be due to competing pathways for methionine between its role as a cofactor for epigenetic modifiers through SAM and in the biosynthesis of high dimethylsulphoniopropionate (DMSP) production in stressed corals^33^. DMSP can act to stabilize osmotic imbalance^34^, which could occur during bleaching due to depletion of intracellular osmolytes such as glycerol and free amino acids under the reduced metabolic state of dysbiosis^35^. SAM also plays critical role as a methyl donor for epigenetic modifier enzymes such as DNA methyltransferases (DNMTs) and histone methyltransferases (HMTs)^36^ that provide regulation of gene expression and expression variability^37^. Therefore, the availability of SAM could further exacerbate energetic demands due to metabolic depression and the cost of bleaching stress response, as regulation of the expression of essential genes in stress response and repair would be further hampered.

**Fig. 3.**
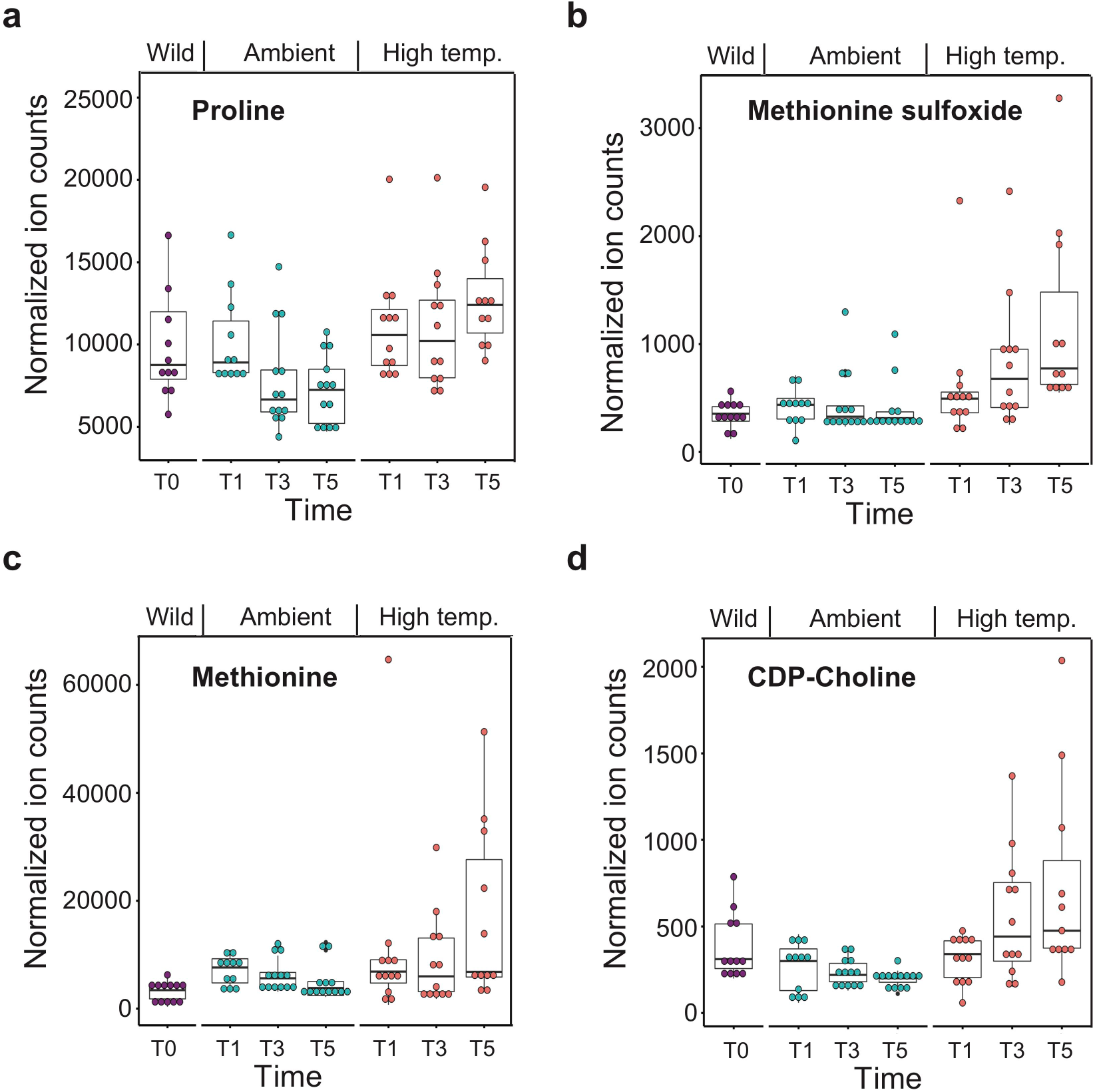
Production of known metabolites by *M. capitata* under thermal stress. (**a**) Accumulation of proline, (**b**) accumulation of methionine sulfoxide, (**c**) accumulation of methionine, and (**d**) accumulation of cytidine diphosphate (CDP)-choline. Differences between the T5 ambient and high temperature groups are significant as determined by a two-tailed Student’s T-Test with *p*-values of 0.03982, 0.00472, 0.04731, and 0.02344 for a-d, respectively.

To gain additional insights into dipeptide function in corals, we generated co-occurrence networks that included the validated and predicted dipeptides with known metabolic features. We tested the idea that dipeptide production is co-regulated and cluster under thermal stress. To address this hypothesis, Louvain communities^38^ in networks were detected and a dipeptide score was computed using the following formula to determine the average dipeptide by community (ADPC) score (Table S2) in which *n*=nodes, *dp*=dipeptides, and *okm*=other known metabolites.

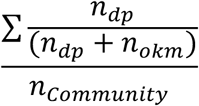

This analysis shows that the ADPC score for dipeptides becomes smaller from T1 to T5, indicative of greater clustering (Fig. 4a). This pattern is apparent in the networks of *M. capitata* at T5 whereby large clusters of dipeptides occur in the thermal stress networks (Fig. S4). Here we focused on two subnetworks in the T5 thermal stress treatment that offer key insights. The first shows that the Montiporic acids A and D (MA-A, and the highly abundant MA-D; Fig. 1d), presumably of host origin, link two key stress responses, carbohydrate metabolism and protective osmolyte production (Fig. 4b). The production of organic osmolytes such as betaine, sorbitol (and its precursor, glucose), sucrose, and trimethylamine N-oxide (TMAO) are all associated with the stress response^39^. These are all co-regulated and increase in abundance in the coral holobiont, with the algal symbiont as the likely source of these compounds^40^. Organic acids are also important intermediates in energy production and as sensing molecules^41^. On the left side of this subnetwork are metabolites involved in carbohydrate metabolism, including pyruvate, that can be fed into the TCA cycle to produce ATPs. Another metabolite, histamine, a neurotransmitter and bioenergetic intermediate that may be involved in injury signaling in *M. capitata*^42^, accumulates from T1 to T5 (Fig. 4c). A second subnetwork (Fig. 4d) shows the co-regulation of many dipeptides, including RQ and other characterized compounds (KQ, RV, RA, RT). In addition to the functions described above as a potential response to oxygen stress, dipeptides and free amino acids (right side of the subnetwork) are important, and readily assimilated sources of nitrogen for the algal symbiont and can be used to generate energy via the TCA cycle or metabolized in the purine pathway^40^. The putative methylation-related metabolites, methionine and methionine-sulfoxide, discussed above are in this subnetwork. A key metabolite involved in phospholipid metabolism, cell signaling, glutamate transport, and an intermediate in betaine synthesis^39^, CDP-choline, also shows steady accumulation in *M. capitata* from T1-T5 (Fig. 3d). In support of our findings, a recent study that focused on metabolites predictive of coral bleaching sensitivity identified betaine as a key marker of resilience in *M. capitata*^43^.

**Fig. 4.**
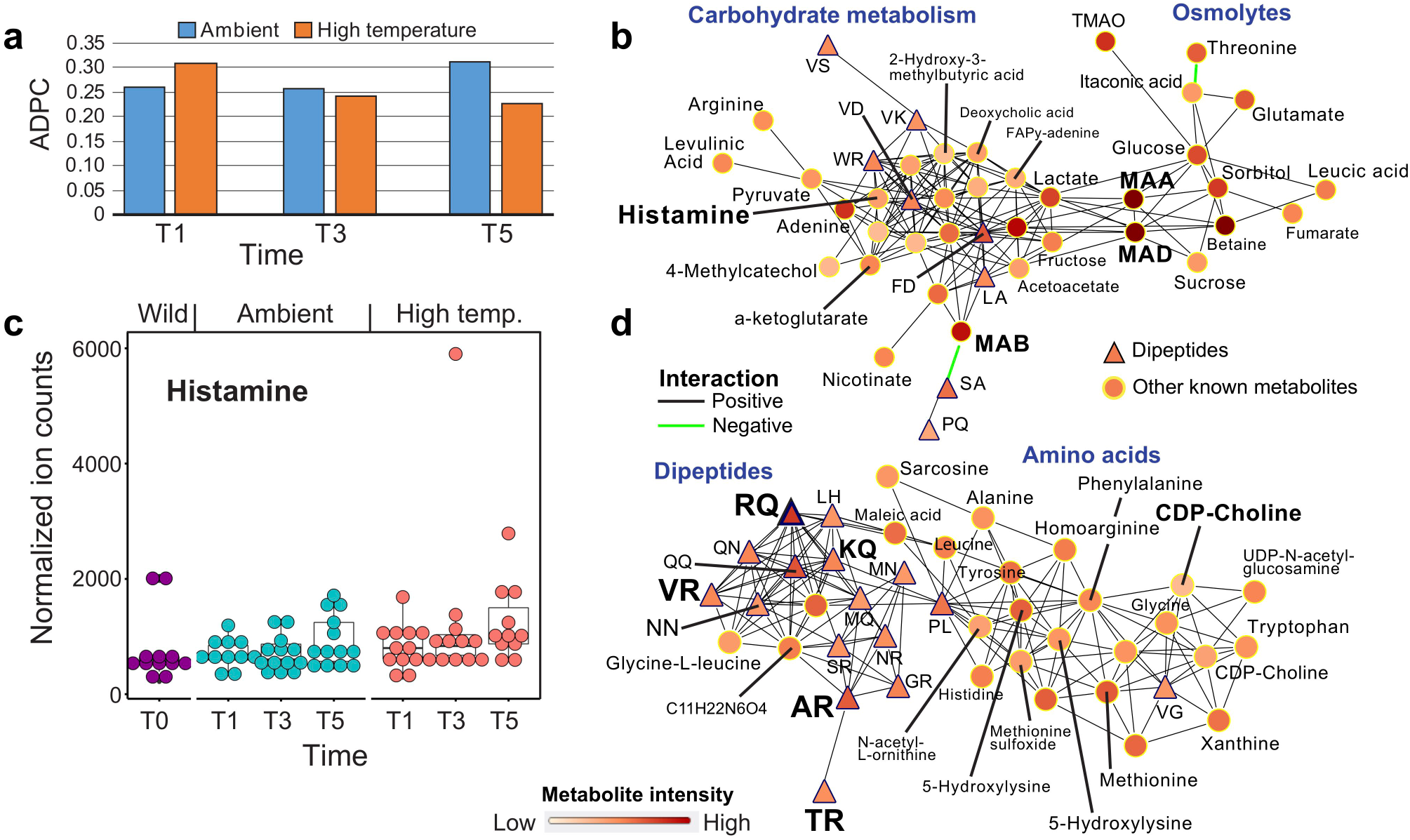
Network analysis of *M. capitata* metabolites. (**a**) ADPC scores for ambient and thermal treatments at T1, T3, and T5. (**b**) Subnetwork at thermal stress T5 showing the relationship between carbohydrate metabolism and osmolyte accumulation. Dipeptides are shown as triangles (under both positive and negative ionization modes) and other metabolites as circles with annotations, when available. Metabolite intensity and type of correlation are shown in the legend. (**c**) Histamine amount as a function of time of thermal stress in lab (T1-T5) and wild colonies. (**d**) Subnetwork at thermal stress T5 showing the relationship between dipeptide and amino acid accumulation.

## Comparison to *Aiptasia*

To investigate the broader distribution of metabolites diagnostic of coral thermal stress, we generated metabolomic data from the sea anemone model *Aiptasia* (Cnidaria). *Aiptasia* is a member of the sister lineage of stony corals^44^ and harbors dinoflagellate symbionts but does not biomineralize^45,46^. Comparison of data gathered from symbiotic and aposymbiotic (alga free, after bleaching [see Methods]) *Aiptasia* individuals showed a significant difference in the accumulation of multiple known and unknown compounds present in corals (Fig. S3b), including RQ and other dipeptides (Fig. S5). Because the experimental design differs between the coral and *Aiptasia* work, we cannot draw a direct link between dipeptides and increasing thermal stress in the latter. However, the increased production of the same dipeptides in symbiotic *Aiptasia* and in thermally stressed *M. capitata* support their role as a response to the presence or absence of algal symbionts. Furthermore, the data suggest that these dipeptides are likely to be generated by the animal host due to their presence in phylogenetically distantly related cnidarian species and occurrence in alga-free *Aiptasia* (Fig. S5). If in fact of animal provenance, then the impact of these (and other) dipeptides on algal metabolism is significant and supports host-regulation of symbiont biology during thermal stress.

## Summary

We used a controlled time course experiment to create a metabolome-based understanding of the coral response to thermal stress in *M. capitata* and *P. acuta*. Using color scores to assess the rate of bleaching, samples were collected at regular intervals for both targeted and untargeted LC-MS analysis under both positive and negative polarities. These results provide the platform to address differences in resilience to thermal stress between coral colonies that are located in close proximity in reefs or are generated using controlled breeding experiments. Elucidating how metabolites associated with DNA and gene modifications events, such as methylation, can alter expression patterns and effect resilience is of importance to conservation efforts. The finding of high accumulation of MAs and their central position in the metabolite networks was surprising because these polyacetylenic carboxylic acids are more abundant than many metabolites in central metabolic pathways such as glycolysis and the TCA cycle. This suggests that coral animals divert a large part of metabolic resources to the production of secondary metabolites. Equally surprising was the critical role of dipeptides, such as RQ, in coral stress biology. By generating additional structures of novel stress-associated coral metabolites and their accumulation in different species and populations, we hope in the future, to understand the intersection of holobiont metabolomes, (epi)genetic architecture of these meta-organisms, and coral resilience in the field.

## Methods

### Cultivation of coral nubbins

From the waters of Kāne’ohe Bay, HI, four colonies of each coral species *Montipora capitata* and *Pocillopra acuta* were identified and collected under SAP 2019-60. Each of the four colonies for each species was fragmented into 30 pieces at the Hawai’i Institute of Marine Biology, located on Moku o Lo’e in Kāne’ohe Bay, HI, and hot-glued to labeled plugs. The 30 glued nubbins of each genotype were then randomly distributed among six tanks (∼32 L, 48.3 × 38.1 ×17.8cm, L x W x H), leaving 5 replicates per genotype in each tank for a total of 40 coral nubbins per tank, and 120 coral nubbins per species. Tanks were placed in a flow through system that had a steady supply of water directly from Kāne’ohe Bay with an average flow rate of (173.8 ± 73 Lh^-1^, mean ± sd). Each tank was fitted with a submersible Pump (Hydor 200 gph), a Hobo Water Temp Pro temperature logger (Onset Computer Corp, operation range = -40° to 70°C, resolution = 0.02°C at 25°, accuracy = ±0.21°C from 0° to 50°C), an APEX temperature probe (Neptune Systems) and two heaters (Aqueon 300W Heater set to 31°C, DaToo 300W Glass Heater set to 34°C). The temperature in the tanks was controlled by powering off and on the heaters based on set points in the Apex aquarium controller (Neptune Systems). Light was set for a 12hr light dark cycle using Arctic-T247 Lights (Ocean Revive).

### Assessment of coral bleaching

Each sample was photographed using a digital camera with a Red/Blue/Green color standard. Red/Blue/Green values were extracted in ImageJ^47^ from the coral were standardized to the color cards by dividing the experimental value observed in the coral against the corresponding actual recorded value from the color card^21^. Using the normalized intensity values from each color channel, a bleaching score was quantified as PC1 from principle components analysis. As stress is prolonged and bleaching becomes more pronounced, the Red/Blue/Green color readings from the coral will equalize around the same number, because white is an equal expression of all colors. All nubbins available at each time points were chosen used for color assessment.

### Experimental design

Once fragmented, the coral nubbins were allowed an acclimation period of 5 days at ambient temperature (26.84 ± 0.50, mean ± sd) before initiating the temperature ramping. Tanks were randomly assigned to treatments groups, with tanks 1, 4, and 6 in the ambient treatment and tanks 2, 3, and 5 in the high temperature treatment. The high temperature treatment tanks were set to increase by ∼0.4°C every 2 days for a total of 9 days (5/13-5/22/19) until they were between 30.5°C to 31.0°C (Figure S1). Treatments of 30.33 ± 0.35 (mean ± sd) for high and 27.67 ± 0.34 (mean ± sd) for ambient were held for the remainder of the experiment, which lasted 16 days (Figure S1). High tanks were on average 2.66°C above ambient when the ramp was completed. Temperature readings from the Hobo loggers were confirmed using spot checks with a handheld digital certified thermometer (Control Company accuracy = ± 0.05 °C, resolution = 0.001°C) ∼2-3 times daily. Light measurements were taken to quantify photosynthetically active radiation (Photosynthetic Photon Flux Density in µmol photons m-2 s-1) using an underwater cosine light sensor and meter (MQ-510: Full Spectrum Underwater Quantum Meter Apogee), and did not differ substantially between tanks (Ambient = 345 ± 25.2 High = 340 ± 29.8 µmol photons m^-2^ s^-1^; mean ± sd). The pH on the total scale was measured with a glass probe (Mettler Toledo InLab Expert Pro pH probe #51343101; accuracy = ±0.2 mV, resolution = 0.1 mV) and handheld meter (Thermo Scientific Orion Star A series A325) and showed no significant differences between the treatments (Ambient = 7.91 ± 0.07, High = 7.91 ± 0.04; mean ± sd; t = 0.30332, df = 78.732, *p*-value = 0.7624).

### Coral Sampling

Three sampling points were selected on 5/22/19 (T1), 6/3/19 (T3), and 6/7/19 (T5) based on reaching maximum treatment temperature (Figure S1), where coral Color Score began to diverge by treatment, and where coral Color Scores were maximized between treatments within species. Corals were sampled at ∼14:30 at each timepoint by removing them from their treatment only touching the plastic bases and inserting them into sterile whirlpaks that were immediately submerged in liquid nitrogen and transferred to -80°C until metabolite extraction.

### Metabolite extraction from coral nubbins

Metabolites were extracted using a protocol optimized for water soluble polar metabolite analysis on LCMS. After the coral nubbins were harvested, placed in zip tie bags, and flash frozen in liquid nitrogen to preserved their metabolic integrity. These were processed for metabolomic analysis immediately or stored at -80°C. The extraction buffer used was a (v/v/v) solution of 40:40:20 (Methanol:Acetonitrile:Water) + 0.5% Formic Acid. The extraction buffer was stored at -20°C prior to usage and immediately preceding the metabolite extraction 1 mL added to a glass 2 mL Dounce homogenizers that had chilled on ice. Pieces of the preserved nubbins were then clipped, weighed, and added to the cold extraction buffer in the Dounce and left to incubate for 5 min. The pestle of the Dounce was then used to homogenize the coral tissue until there was a visible accumulation of coral skeleton at the bottom of the Dounce and the homogenate was visibly pigmented. An additional 500 µL aliquot of cold 40:40:20 + 0.5% formic acid extraction buffer was then used to rinse down the sides of the Dounce and pestle. The total 1.5 mL volume was then strained through a 100 µm Cell Strainer into a 50 mL receptacle. There was a visible amount of skeleton collected in the strainer. The rest of the homogenate was then transferred to a 1.5 mL Eppendorf tube and vortexed for 10 sec and then centrifuged for 10 min at highest speed at 4°C. After centrifugation, there was a pellet at the bottom of the tube. A final 500 µL aliquot of the homogenate was then pipetted to a second clean Eppendorf tube, to which 44 µL of 15% NH_4_HCO_3_ was added to neutralize the acid in the buffer. This was the final extract and ready to be loaded to instrument vials for analysis.

### Cultivation of *Aiptasia*

All animals were from the clonal population CC7, which naturally contains the algal symbionts of the Symbiodinuaceae clade A species *Symbiodinium linuchae*^,48,49^. CC7 animals were made aposymbiotic through repeated cycles of cold shock with addition of artificial sea water at 4°C and subsequent incubation at 4°C for 4 hours, followed by a 1-2-day photosynthesis inhibitor diuron treatment (Sigma-Aldich) at 50 µM^50,51^. This resulted in the strain CC7-Apo. CC7-Apo animals subsequently exposed to algae of the clonal axenic strain SSB01 Symbiodinuaceae clade B species *Breviolum minutum* and grown under standard conditions until the algal populations reached steady state. The CC7-SSB01 strain was periodically assessed by sequencing PCR-amplified fragments of cps23S (chloroplast rDNA), 18S (nuclear rDNA), and/ or ITS2 (nuclear rDNA), to ensure that repopulation with *Symbiodinium linuchae* or any other algal symbiont type had not occurred. CC7-Apo and CC7-SSB01 were used for metabolomic analysis. *Aiptasia* were aliquoted individually to 1.5 mL Eppendorf tubes in 500µL of artificial seawater, flash frozen, and stored at -80°C until processing at Rutgers University.

### Metabolite extraction from *Aiptasia*

Metabolites were extracted using a protocol optimized for water soluble polar metabolite analysis on LCMS. The extraction buffer used was a (v/v/v) solution of 40:40:20 (Methanol:Acetonitrile:Water) + 0.5% Formic Acid and was stored at -20°C prior to usage. CC7-Apo and CC7-SSB01 were pooled together intragroup-wise together to produce six replicates each having an input weight of 25 mg in a 1.5 mL Eppendorf tube. Replicates were kept on dry ice prior to extraction. A total of 500 µL of extraction buffer was added, then samples were vortexed for 10 sec and transferred to crushed ice to incubate for 10 min. Samples were then centrifuged for 10 min and the supernatant was then transferred to a 1.5 mL Eppendorf tube and the procedure repeated for a second round of extraction. A total of 1 mL of supernatant was then centrifuged and a final 500 µL aliquot of the homogenate was then pipetted to a clean Eppendorf tube, to which 44 µL of 15% NH_4_HCO_3_ was added to neutralize the acid in the buffer. This was the final extract and ready to be loaded to instrument vials for analysis.

### UHPLC chromatography conditions

The HILIC separation was performed on a Vanquish Horizon UHPLC system (Thermo Fisher Scientific, Waltham, MA) with XBridge BEH Amide column (150 mm × 2.1 mm, 2.5 µm particle size, Waters, Milford, MA) using a gradient of solvent A (95%:5% H_2_O:acetonitrile with 20 mM acetic acid, 40 mM ammonium hydroxide, pH 9.4), and solvent B (20%:80% H_2_O:acetonitrile with 20 mM acetic acid, 40 mM ammonium hydroxide, pH 9.4). The gradient was 0 min, 100% B; 3 min, 100% B; 3.2 min, 90% B; 6.2 min, 90% B; 6.5 min, 80% B; 10.5 min, 80% B; 10.7 min, 70% B; 13.5 min, 70% B; 13.7 min, 45% B; 16 min, 45% B; 16.5 min, 100% B and 22 min, 100% B. The flow rate was 300 µl/min. Injection volume was 5 µL and column temperature was 25 °C. The autosampler temperature was set to 4°C and the injection volume was 5µL.

### Full scan mass spectrometry

The full scan mass spectrometry analysis was performed on a Thermo Q Exactive PLUS with a HESI source which was set to a spray voltage of -2.7 kV under negative mode and 3.5 kV under positive mode. The sheath, auxiliary, and sweep gas flow rates of 40, 10, and 2 (arbitrary unit) respectively. The capillary temperature was set to 300°C and aux gas heater was 360°C. The S-lens RF level was 45. The m/z scan range was set to 72 to 1000 m/z under both positive and negative ionization mode. The AGC target was set to 3e6 and the maximum IT was 200ms. The resolution was set to 70,000. The MS1 data were processed using MAVEN^52^.

### Parallel reaction monitoring mass spectrometry

The MS^2^ spectra generation was performed on a Thermo Q Exactive PLUS with a HESI source which was set to a spray voltage of -2.7kV under negative mode and 3.5kV under positive mode. The sheath, auxiliary, and sweep gas flow rates of 40, 10, and 2 (arbitrary unit) respectively. The capillary temperature was set to 300°C and aux gas heater was 360°C. The S-lens RF level was 45. The m/z scan range was specified at m/z 301.16298 under negative ionization mode and m/z 303.17753 under positive ionization mode and monitored for the full 22 min run time. The AGC target was set to 2e5 and the maximum IT was 100 ms. The resolution was set to 70,000. The Isolation window was set to 2.0 m/z. Collision energy was set to a stepwise 30,50,80 NCE. Our results suggest a single MS^2^ spectrum may contain irrelevant m/z signals and the pseudospectrum, generated by correlating multiple MS^2^ spectra to the MS^1^ extracted ion chromatogram is an effective approach to “clean” the MS^2^ spectrum.

### Synthesis of standards

The dipeptide standards were synthesized at 95% purity by GeneScript USA (Piscataway, New Jersey). All standards were produced in quintuplet, shipped at 25°C, and stored at -20°C until prepared for analysis on the mass spectrometer. Parallel reaction monitoring was used to generate MS^2^ spectra for comparison with samples.

### Network analysis

The R-package DGCA^53^ was used to determine the correlation between pairs of metabolites respectively under ambient and stressed conditions for *M. capitata*. The pairwise correlation was calculated with the function *matCorr* using the Pearson method. The functions *matCorSig* and *adjustPVals*, respectively, used to calculate and adjust (with the Benjamini-Hochberg method) the correlations *p*-values. Only pairs with an adjusted *p*-value ≤ 0.05 were retained to construct the co-occurrence networks.

## Data availability

Raw experimental data and analytical code are available at https://github.com/hputnam/Coral_Hospital/releases/tag/v1 and will be archived at BCO-DMO upon acceptance for publication.

## Acknowledgments

This work was supported by a seed grant awarded to D.B. from the office of the Vice Chancellor for Research and Innovation at Rutgers University and by NSF grants NSF-OCE 1756616 and NSF-OCE 1756623 (D.B., H.M.P.). D.B. was supported by a NIFA-USDA Hatch grant (NJ01170).

## Author contributions

D.B. and X.S. conceived the project; A.W. and E.C.N. performed the bioinformatic analyses; J.S.P. performed the network analysis; H.M.P. helped design the experiments and provided coral tissues for analysis; H.M.P. and D.C. conducted the coral experiment; P.A.C. provided *Aiptasia* tissues; all authors provided feedback on the manuscript and experimental design. A.W., E.N.C., X.S. and D.B. wrote the article with the help from all authors.

## Supplements

**Fig. S1. A) Color scores for the B) ambient and high temperature treated coral species *Montipora capitata* and *Pocillopora acuta* at the Hawai**’**i Institute of Marine Biology (HIMB).** Ambient temperature tanks are shown in variations of red and high temperature tanks n variations of blue. The color scores represent a proxy for algal symbiont density in coral holobionts with low values indicating bleaching phenotype. The sharp score decrease for *P. acuta* under ambient tank conditions is explained by the unexpected warming event that occurred in Kāne’ohe Bay, O’ahu from which the culture water was drawn. Vertical grey lines indicate sampling points T1 (5/22/19), T3 (6/03/19), and T5 (6/7/19).

**Fig. S2. Metabolomic analysis of Hawaiian corals.** (**a**) Accumulation pattern of metabolites in *M. capitata* at different significance levels from T1-T5 under ambient and thermal stress conditions. (**b**) Montiporic acids (MAs) identified in the coral holobiont. (**c**) Accumulation of different MAs in *M. capitata* (MC) and *P. acuta* (PC) in samples collected from the field.

**Fig. S3. Volcano plots of metabolites generated from *P. acuta* and *Aiptasia* under positive ionization mode.** (**a**) Analysis of *P. acuta* data under ambient and high temperature treatments at the three sampling points. (**b**) Analysis of *Aiptasia* under symbiotic and aposymbiotic (alga free) conditions. Putative dipeptides are shown with the filled brown circles in all plots.

**Fig. S4. Annotated network of metabolite co-occurrence in *M. capitata* at T5.** (**a**) Ambient treatment network and (**b**) thermal stress network. Dipeptides are shown as triangles (under both positive and negative ionization modes) and other metabolites as circles with annotations, when available. Metabolite intensity and type of correlation are shown in the legend.

**Fig. S5. Analysis of metabolites in *Aiptasia*.** (**a**) Metabolite (RQ) amount in symbiotic and aposymbiotic (bleached) individuals of *Aiptasia*. (**b**) Metabolite (KQ) amount in symbiotic and aposymbiotic individuals of *Aiptasia*. (**c**) Metabolite (RV) mount in symbiotic and aposymbiotic individuals of *Aiptasia*. (**d**) Metabolite (RA) amount in symbiotic and aposymbiotic individuals of *Aiptasia*.

## Tables

**Table S1**. Coral peptide identification by retention time and MS^2^ spectra matching.

**Table S2**. ADPC scores for dipeptides in the networks derived from *M. capitata* thermal stress experiments.

