## Supplemental figures and tables for "Metabolome shift associated with thermal stress in coral holobionts"

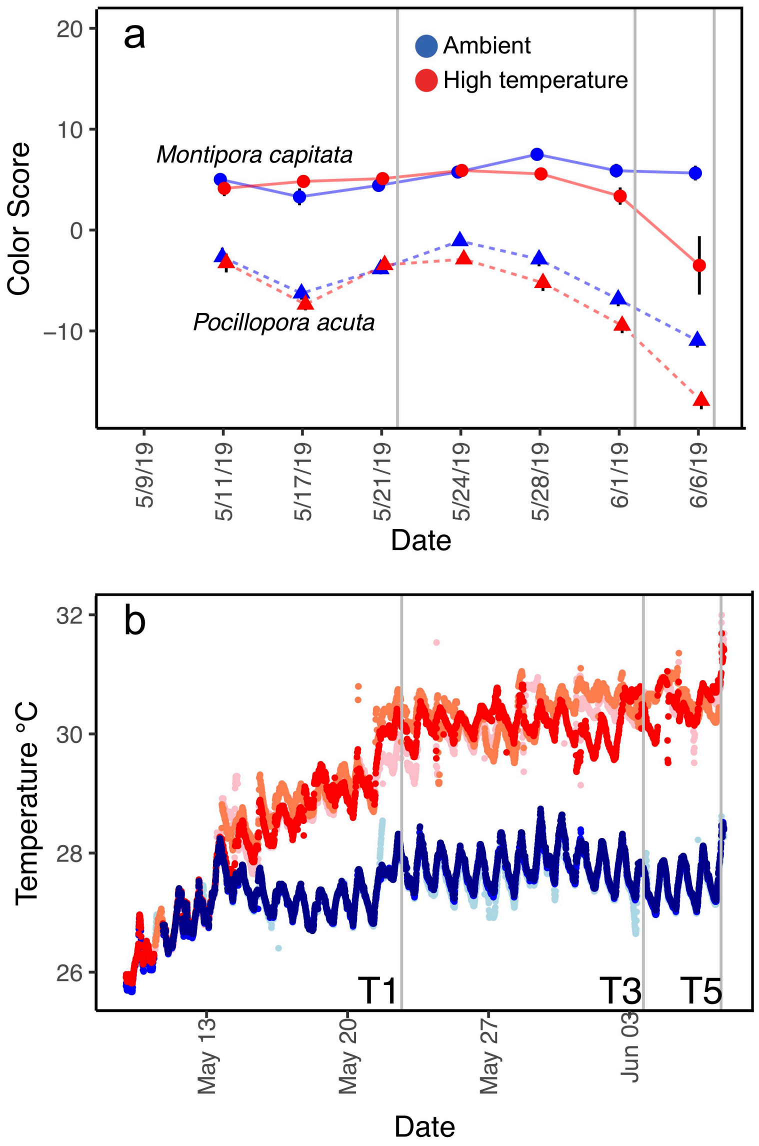

a
Ambient
High temperature
Montipora capitata
Pocillopora acuta
Color Score
20
10
0
−10
5/9/19
5/11/19
5/17/19
5/21/19
5/24/19
5/28/19
6/1/19
6/6/19
Date
b
Temperature °C
32
30
28
26
T1
T3
T5
May 13
May 20
May 27
Jun 03
Date

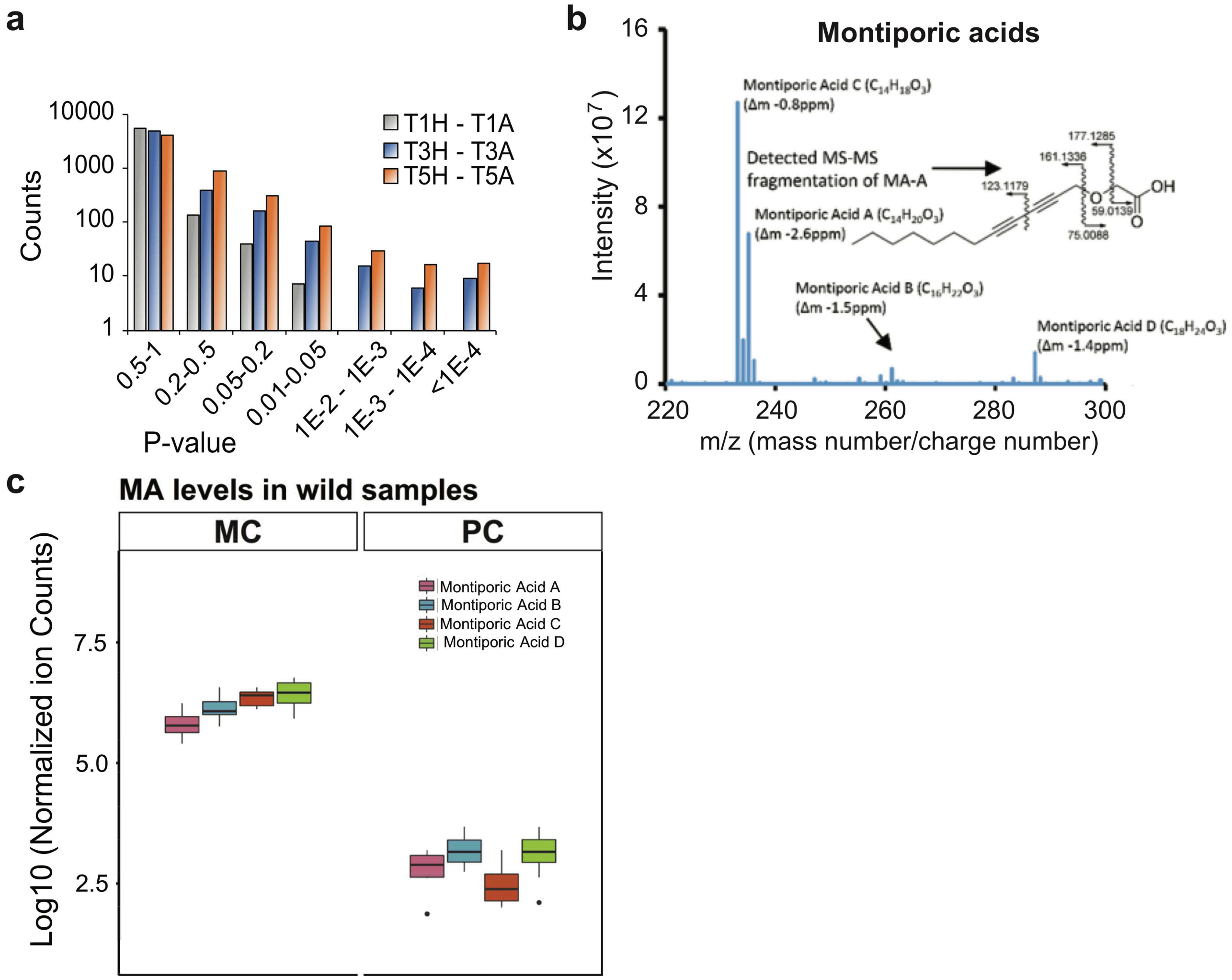
a
10000
1000
100
10
1
Counts
T1H - T1A
T3H - T3A
T5H - T5A
0.5-1
0.2-0.5
0.05-0.2
0.01-0.05
1E-2 - 1E-3
1E-3 - 1E-4
<1E-4
P-value
b
Montiporic acids
16
12
8
4
0
Intensity (x10^7)
Montiporic Acid C (C14H18O3)
(Δm -0.8ppm)
Detected MS-MS
fragmentation of MA-A
Montiporic Acid A (C14H20O3)
(Δm -2.6ppm)
177.1285
161.1336
123.1179
59.0139
75.0088
OH
O
Montiporic Acid B (C16H22O3)
(Δm -1.5ppm)
Montiporic Acid D (C18H24O3)
(Δm -1.4ppm)
220
240
260
280
300
m/z (mass number/charge number)
c
MA levels in wild samples
MC
PC
Montiporic Acid A
Montiporic Acid B
Montiporic Acid C
Montiporic Acid D
7.5
5.0
2.5
Log10 (Normalized ion Counts)

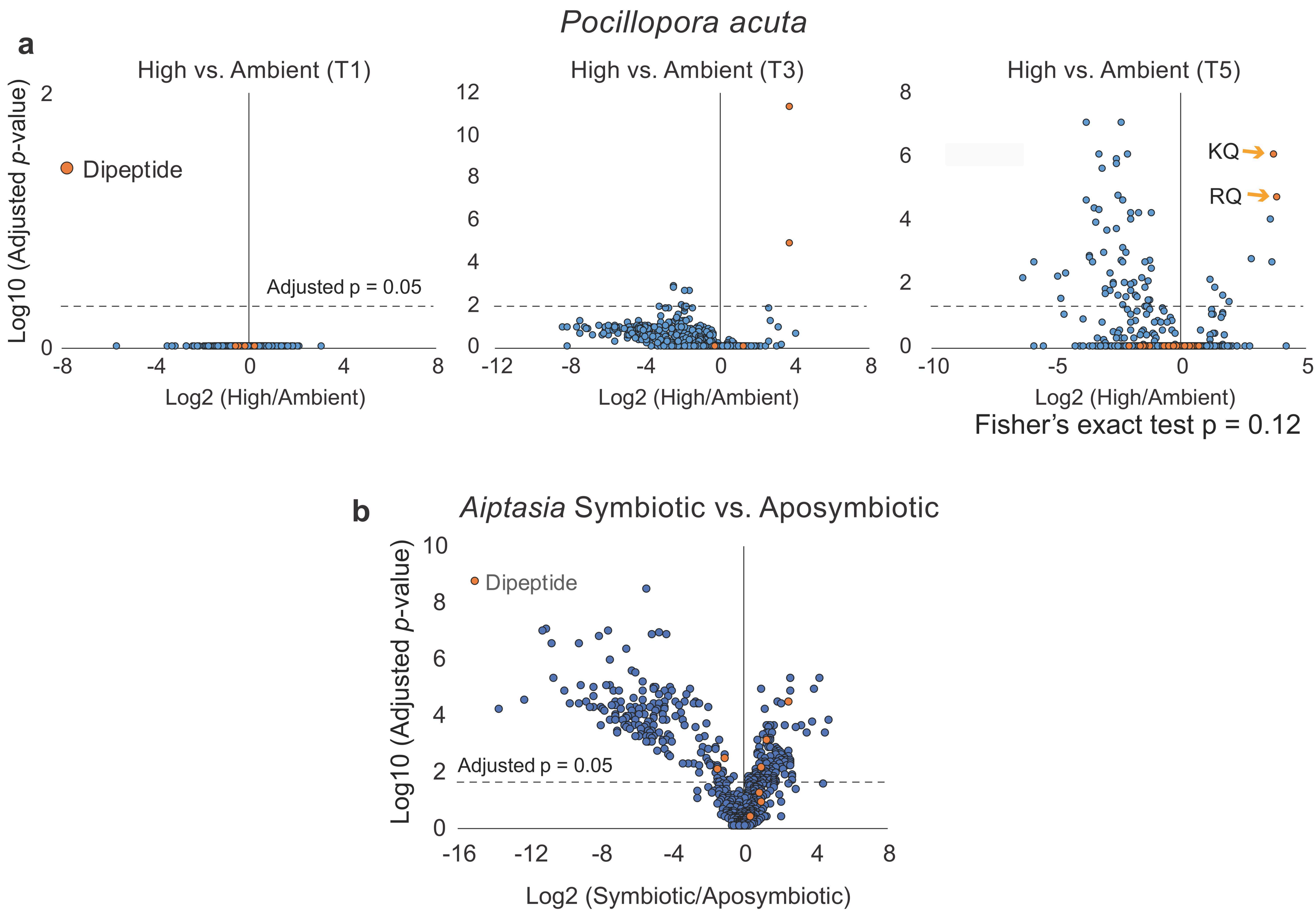
Pocillopora acuta
a
High vs. Ambient (T1)
Dipeptide
Adjusted p = 0.05
Log10 (Adjusted p-value)
Log2 (High/Ambient)
High vs. Ambient (T3)
Log2 (High/Ambient)
High vs. Ambient (T5)
KQ
RQ
Log2 (High/Ambient)
Fisher's exact test p = 0.12
b
Aiptasia Symbiotic vs. Aposymbiotic
Dipeptide
Adjusted p = 0.05
Log10 (Adjusted p-value)
Log2 (Symbiotic/Aposymbiotic)

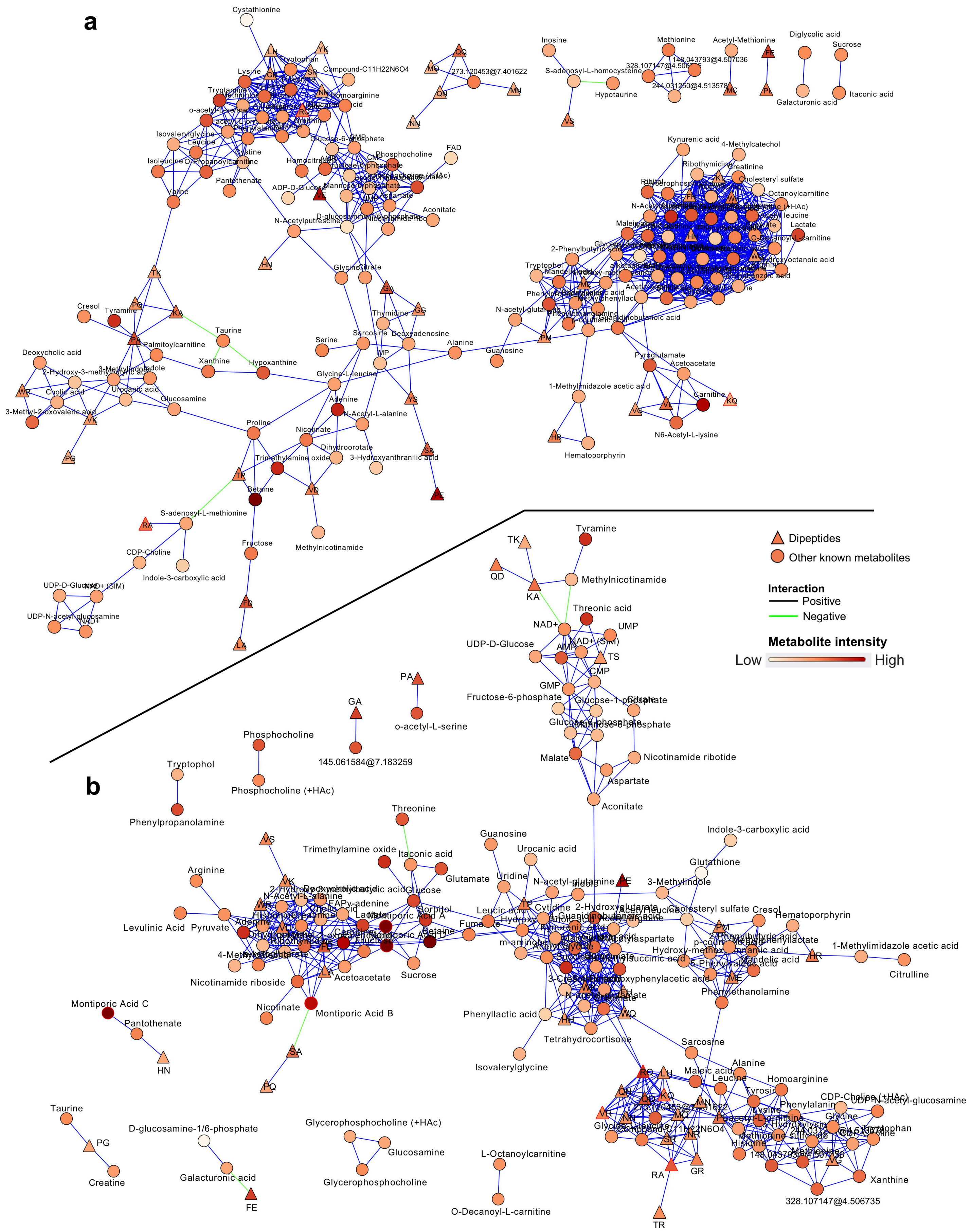

a
b
Dipeptides
Other known metabolites
Interaction
Positive
Negative
Metabolite intensity
Low
High
Cystathionine
Inosine
Methionine
Acetyl-Methionine
Diglycolic acid
Sucrose
Compound-C11H22N6O4
273.120453@7.401622
S-adenosyl-L-homocysteine
Hypotaurine
244.031250@4.513578
Galacturonic acid
Itaconic acid
Kynurenic acid
4-Methylcatechol
Ribothymidine
Creatinine
Cholesteryl sulfate
Octanoylcarnitine
Lactate
Isovalerylglycine
Leucine
Isoleucine
Cystine
Valine
Pantothenate
Phosphocholine
FAD
ADP-D-Glucose
Aconitate
N-Acetylputrescine
Homoarginine
Glycine
Citrate
Thymidine
Tryptophol
2-Phenylbutyric acid
N-acetyl-glutamine
Cresol
Tyramine
Taurine
Palmitoylcarnitine
Xanthine
Hypoxanthine
Serine
Sarcosine
Deoxyadenosine
Alanine
Guanosine
Deoxycholic acid
Cholic acid
Urocanic acid
Glucosamine
Glycine-L-leucine
Pyroglutamate
Acetoacetate
1-Methylimidazole acetic acid
Carnitine
N6-Acetyl-L-lysine
Hematoporphyrin
3-Methyl-2-oxovaleric acid
Proline
Adenine
N-Acetyl-L-alanine
Nicotinate
Dihydroorotate
3-Hydroxyanthranilic acid
Trimethylamine oxide
Betaine
S-adenosyl-L-methionine
Methylnicotinamide
Fructose
CDP-Choline
Indole-3-carboxylic acid
UDP-D-Glucose
NAD+ (SIM)
UDP-N-acetyl-glucosamine
NAD+
Tyramine
Methylnicotinamide
Threonic acid
UMP
UDP-D-Glucose
AMP
CMP
GMP
Fructose-6-phosphate
Glucose-1-phosphate
Mannose-6-phosphate
Malate
Nicotinamide ribotide
Aspartate
Aconitate
o-acetyl-L-serine
Phosphocholine
145.061584@7.183259
Phosphocholine (+HAc)
Tryptophol
Phenylpropanolamine
Threonine
Guanosine
Indole-3-carboxylic acid
Trimethylamine oxide
Itaconic acid
Urocanic acid
Glutathione
Glutamate
Uridine
N-acetyl-glutamine
3-Methylindole
Arginine
Glucose
Sorbitol
Levulinic Acid
Pyruvate
Montiporic Acid A
Leucic acid
Cytidine
2-Hydroxyglutarate
Cholesteryl sulfate
Cresol
Hematoporphyrin
Fumarate
Kynurenic acid
Acetylaspartate
1-Methylimidazole acetic acid
Citrulline
Acetoacetate
Sucrose
Nicotinamide riboside
Mandelic acid
Phenylethanolamine
Montiporic Acid C
Nicotinate
Montiporic Acid B
Phenyllactic acid
Pantothenate
Tetrahydrocortisone
Sarcosine
Isovalerylglycine
Alanine
Maleic acid
Homoarginine
Leucine
Tyrosine
Phenylalanine
CDP-Choline (+HAc)
UDP-N-acetyl-glucosamine
Taurine
D-glucosamine-1/6-phosphate
Glycerophosphocholine (+HAc)
Glucosamine
L-Octanoylcarnitine
Creatine
Galacturonic acid
Glycerophosphocholine
O-Decanoyl-L-carnitine
Xanthine
328.107147@4.506735

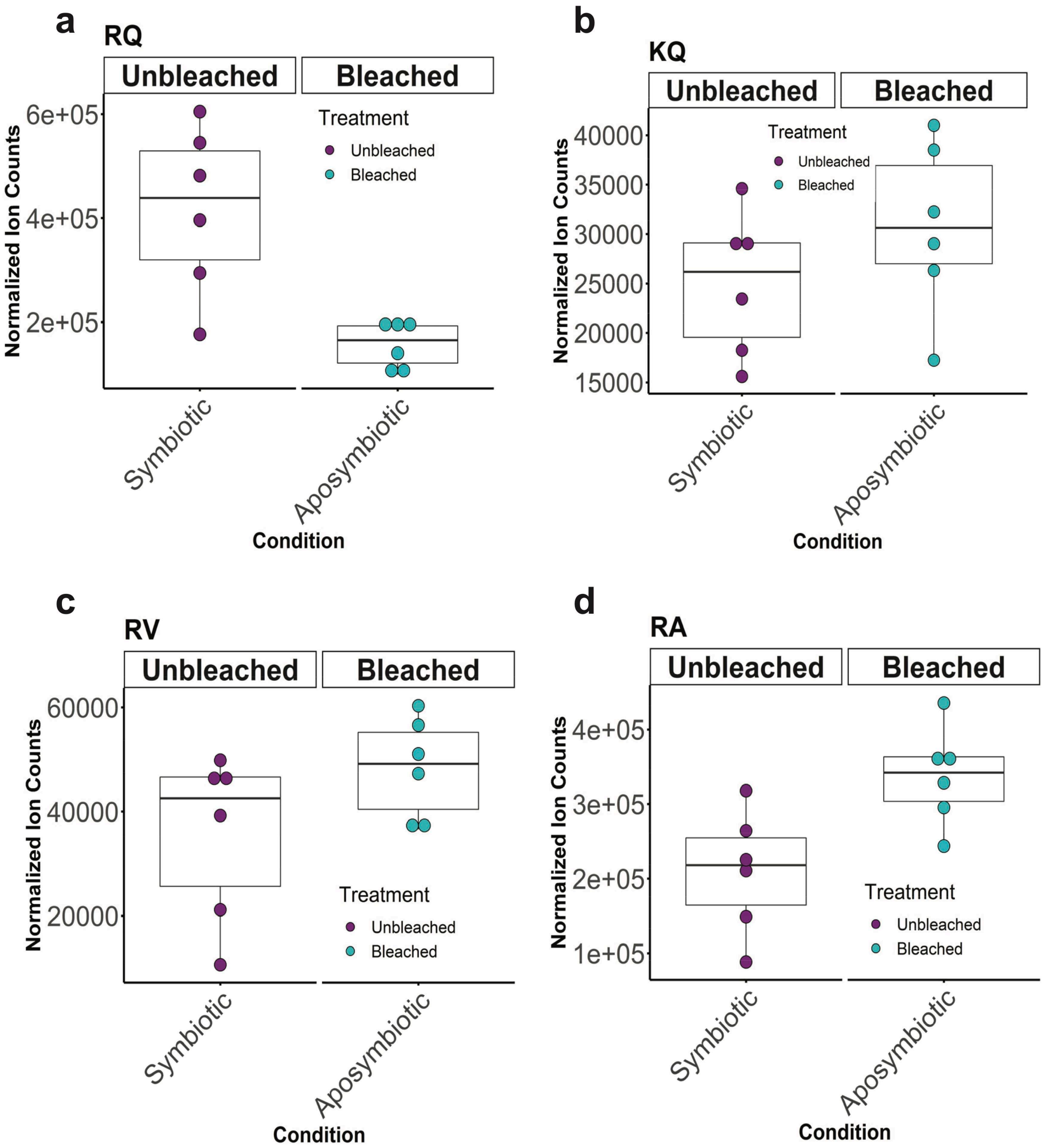

a
RQ
Unbleached
Bleached
Treatment
Unbleached
Bleached
Normalized Ion Counts
6e+05
4e+05
2e+05
Symbiotic
Aposymbiotic
Condition
b
KQ
Unbleached
Bleached
Treatment
Unbleached
Bleached
Normalized Ion Counts
40000
35000
30000
25000
20000
15000
Symbiotic
Aposymbiotic
Condition
c
RV
Unbleached
Bleached
Treatment
Unbleached
Bleached
Normalized Ion Counts
60000
40000
20000
Symbiotic
Aposymbiotic
Condition
d
RA
Unbleached
Bleached
Treatment
Unbleached
Bleached
Normalized Ion Counts
4e+05
3e+05
2e+05
1e+05
Symbiotic
Aposymbiotic
Condition

**Table S1**. Coral peptide identification by retention time and $MS^2$ spectra matching.

| Coral Dipeptide | Peptide Standard | RT (min) | ΔRT (min) | $MS^2$ Spectra Matching |
|---|---|---|---|---|
| **$C_{11}H_{22}N_6O_4$ (RT 9.05min)** | **RQ** | **9.1** | **0.05** | **99.69%** |
| | QR | 8.82 | -0.23 | 85.33% |
| | ER-NH2 | 9.38 | 0.33 | 77.86% |
| | RE-NH2 | 9.3 | 0.25 | 97.36% |
| | γ-ER-NH2 | 10.84 | 1.79 | 86.41% |
| | AGR | 8.71 | -0.34 | 86.95% |
| | RAG | 8.62 | -0.43 | 96.64% |
| | RGA | 8.62 | -0.43 | 96.83% |
| **$C_{11}H_{22}N_4O_4$ (RT 9.96min)** | **KQ** | **9.98** | **0.02** | **99.15%** |
| | QK | 9.54 | -0.42 | 82.35% |
| | EK-NH2 | 10.21 | 0.25 | 91.62% |
| | KE-NH2 | 10.11 | 0.15 | 99.46% |
| | γ-EK-NH2 | 12.89 | 2.93 | 98.23% |
| **$C_{11}H_{23}N_5O_3$ (RT 7.64min)** | **RV** | **7.62** | **-0.02** | **98.12%** |
| | VR | 6.81 | -0.83 | 54.50% |
| **$C_9H_{19}N_5O_3$ (RT 8.97min)** | **RA** | **8.95** | **-0.02** | **99.36%** |
| | AR | 8.98 | 0.01 | 86.19% |

**Table S2**. Average dipeptide by community (ADPC) scores for dipeptides in the networks derived from the *M. capitata* lab manipulations. AMB is the ambient condition and HIT is the thermal stress treatment.

| Conditions | Time | ADPC | #Community | DP | OKM | #Nodes | #Edges | Density | Centralization |
|---|---|---|---|---|---|---|---|---|---|
| **AMB** | T1 | 0.2599559 | 25 | 34 | 121 | 155 | 309 | 0.0259 | 0.066 |
| **AMB** | T3 | 0.2573268 | 11 | 45 | 170 | 215 | 2131 | 0.0926 | 0.156 |
| **AMB** | T5 | 0.3119065 | 15 | 45 | 156 | 201 | 933 | 0.0464 | 0.135 |
| **HIT** | T1 | 0.3071103 | 14 | 46 | 160 | 206 | 1012 | 0.0479 | 0.099 |
| **HIT** | T3 | 0.2425315 | 18 | 42 | 147 | 189 | 571 | 0.0321 | 0.069 |
| **HIT** | T5 | 0.2251791 | 16 | 42 | 138 | 180 | 639 | 0.0396 | 0.072 |
